# Dynamic oncogene activity metabolically reprograms the tryptophan– kynurenine–AHR axis driving immune evasion and tumor progression in Ewing sarcoma

**DOI:** 10.1101/2025.05.16.654502

**Authors:** Martha J. Carreño-Gonzalez, Kimberley M. Hanssen, Ahmed Sadik, Alessa L. Henneberg, Maximilian M. L. Knott, Anna C. Ehlers, A. Katharina Ceranski, Zuzanna A. Kolodynska, Malenka Zimmermann, Roland Imle, Anneliene H. Jonker, Florian H. Geyer, Alina Ritter, Ana Banito, Clémence Henon, Olivier Delattre, Didier Surdez, Shunya Ohmura, Florencia Cidre-Aranaz, Christiane A. Opitz, Thomas G. P. Grünewald

**Affiliations:** Hopp Children’s Cancer Center (KiTZ), Heidelberg, Germany; Division of Translational Pediatric Sarcoma Research, German Cancer Research Center (DKFZ), German Cancer Consortium (DKTK), Heidelberg, Germany; National Center for Tumor Diseases (NCT), NCT Heidelberg, a partnership between DKFZ and Heidelberg University Hospital, Germany; Medical Faculty, University of Heidelberg, Heidelberg, Germany; Division of Metabolic Crosstalk in Cancer, German Cancer Research Center (DKFZ), German Cancer Consortium (DKTK), Heidelberg, Germany; Soft-Tissue Sarcoma Junior Research Group, DKFZ, Heidelberg, Germany; Health Technology and Services Research (HTSR) group, University of Twente, Enschede, the Netherlands; SIREDO: Care, Innovation and Research for Children, Adolescents and Young Adults with Cancer, Institut Curie, Paris, France, France; Bone Sarcoma Research Laboratory, Balgrist University Hospital, Zürich, Switzerland; Institute of Pathology, Heidelberg University Hospital, Heidelberg, Germany

**Author notes:** **Address for correspondence** Thomas G. P. Grünewald, MD, PhD Division of Translational Pediatric Sarcoma Research German Cancer Research Center (DKFZ) Im Neuenheimer Feld 280 69120 Heidelberg, Germany.

**Keywords:** Ewing sarcoma, AHR, Kynurenine, EWSR1::ETS, NK cells

## Abstract

The extent to which dynamic changes in oncogene activity shape cancer cell metabolism and drive disease progression remains poorly understood. Ewing sarcoma (EwS), driven by EWSR1::ETS fusion transcription factors, constitutes an ideal model to interrogate this question, as fluctuations in fusion activity direct divergent transcriptional programs. While EWSR1::ETS-high cells display a rather sessile but proliferative phenotype, EWSR1::ETS-low cells are more invasive. Yet, the mechanisms underlying these different phenotypes remain poorly characterized.

Here, by employing an integrative functional metabolomics approach, we link reduced EWSR1::ETS activity in primary EwS tumors to adverse clinical outcome and pronounced activation of the aryl hydrocarbon receptor (AHR) pathway. Low EWSR1::ETS states foster tryptophan catabolism and accumulation of the AHR agonist kynurenine, which in turn promotes an immunosuppressive tumor microenvironment characterized by impaired natural killer (NK) cell cytotoxicity and enrichment of immunoregulatory infiltrates. Functionally, AHR silencing restores NK cell-mediated tumor recognition, while also directly suppressing EwS cell proliferation, clonogenicity, and spheroid growth in plasma-like media. Genetic inhibition of AHR reduces tumor burden and metastatic competence in xenograft models. These findings reveal a mechanistic link between oncogene fluctuation, amino acid metabolism, and immune evasion, positioning AHR as a central mediator of EwS progression and a tractable therapeutic vulnerability.

## INTRODUCTION

Ewing sarcoma (EwS) is the second most common bone and soft tissue tumor in children, adolescents, and young adults (Grünewald *et al*., 2018). Despite intensive multimodal therapy, EwS remains a highly aggressive cancer, with 20–25% of patients presenting with metastatic disease at diagnosis (Grünewald *et al*., 2018; Cidre-Aranaz *et al*., 2022), and recurrent or relapsed disease is associated with dismal survival rates below 30% (Stahl *et al*., 2011; Grünewald *et al*., 2018). While localized tumors can be controlled in the short term, approximately one third of patients relapse, underscoring the urgent need for new therapeutic strategies that address the biological drivers of metastatic spread and treatment resistance, including immune evasion (Grünewald *et al*., 2020; Evdokimova *et al*., 2023).

EwS tumors are driven by a single genetic event – a chromosomal translocation that results in the fusion of a *FET* family gene (*FUS*, *EWSR1*, *TAF15*) with an ETS family transcription factor (mainly *FLI1* or *ERG*). The resulting FET::ETS fusion oncoproteins, most commonly EWSR1::FLI1 (∼85% of cases) and EWSR1::ERG (∼10% of cases) (Delattre *et al*., 1992, 1994; Sorensen *et al*., 1994), act as aberrant transcription factors deregulating the cellular epigenome and transcriptome with tumorigenic implications (May *et al*., 1993; Tomazou *et al*., 2015; Orth *et al*., 2022). Several studies have revealed striking cell-to-cell heterogeneity in EWSR1::ETS expression levels, with high expression favoring proliferation and low expression conferring migratory and invasive traits linked to metastasis (Chaturvedi *et al*., 2012, 2014; Wiles *et al*., 2013; Franzetti *et al*., 2017; Sannino *et al*., 2017; Aynaud *et al*., 2020; Adane *et al*., 2021; Surdez *et al*., 2021). This plasticity suggests that this fluctuating fusion oncoprotein activity may coordinate both tumor growth and dissemination.

Like many sarcomas, EwS tumors are immunologically ‘cold’, characterized by sparse infiltration of cytotoxic immune cells and an enrichment of immunosuppressive myeloid populations (Evdokimova *et al*., 2023). While the degree of CD8 T cell infiltration and immunosuppressive macrophage infiltration has been found to negatively influence EwS patient outcome (Fujiwara *et al*., 2011; Stahl *et al*., 2011), immunotherapies, such as nivolumab, pembrolizumab, and cixutumumab, have thus far shown limited success in early clinical trials for EwS (Malempati *et al*., 2012; Tawbi *et al*., 2017; Davis *et al*., 2020; Palmerini *et al*., 2024). A variety of factors has been attributed to these failures, including low immunogenicity of EwS tumors, MHC class I downregulation, presence of immunosuppressive cells (MDSCs, M2-like tumor associated macrophages, regulatory T cells), and low infiltration of cytotoxic T and NK cells (Malempati *et al*., 2012; Tawbi *et al*., 2017; Davis *et al*., 2020; Evdokimova *et al*., 2023). Understanding how the fusion oncoproteins and their downstream pathways shape the immune landscape therefore represents a critical step toward improving immunotherapeutic opportunities. One emerging axis of interest is metabolic control of immunity. Fusion-driven transcriptomic programs in EwS regulate numerous metabolic genes (Sen *et al*., 2018; Issaq *et al*., 2020; Orth *et al*., 2022), including enzymes of the tryptophan (Trp)–kynurenine (Kyn) pathway (Mutz *et al*., 2016). Metabolites such as Kyn and kynurenic acid (KynA) can activate the aryl hydrocarbon receptor (AHR) (DiNatale *et al*., 2010; Opitz *et al*., 2011), a ligand-activated transcription factor increasingly recognized as a central modulator of tumor immune evasion (Gutiérrez-Vázquez and Quintana, 2018; Rothhammer and Quintana, 2019; Opitz *et al*., 2023).

The functional significance of AHR activation for the EwS oncophenotype and the potential role of the Trp–Kyn–AHR axis in mediating the EwS immune environment, however, is unknown.

Here, we investigate the contribution of the Trp–Kyn–AHR axis to EwS biology and tumor– immune interactions. By integrating metabolomic screens, patient data, and functional experiments, we uncover that AHR signaling is enriched in EwS tumors with poor clinical outcomes, activated in states of low EWSR1::ETS expression, and promotes resistance to NK cell cytotoxicity. Inhibition of AHR restores NK cell-mediated killing, reduces EwS clonogenicity, and abrogates tumor growth in vivo. These findings highlight AHR activation as a fusion-linked vulnerability and nominate the Trp–Kyn–AHR pathway as a tractable target for immunometabolic therapy in EwS.

## RESULTS

### Low EWSR1::ETS activity induces AHR signaling and confers poor overall survival in EwS

Accumulating evidence in cell line models suggests that low EWSR1::ETS activity is linked with a more invasive phenotype (Chaturvedi *et al*., 2012, 2014; Wiles *et al*., 2013; Franzetti *et al*., 2017; Sannino *et al*., 2017; Aynaud *et al*., 2020; Adane *et al*., 2021; Surdez *et al*., 2021). However, whether this translates to a clinical phenotype has – to the best of our knowledge – not been demonstrated to date. To address this important question, we took advantage of a patient cohort with matched transcriptome data and clinical information (n=166 EwS patients) (Musa *et al*., 2019) and assessed the transcriptomic enrichment of a curated EWSR1::ETS activity score (IC^EWS^ score) (Aynaud *et al*., 2020) in these patient samples. K-means clustering of the EWSR1::ETS activity scores separated patients into two groups, referred to as EWSR1::ETS-high and EWSR1::ETS-low. Strikingly, when examining whether these transcriptional groups showed prognostic differences, patients with low EWSR1::ETS activity had significantly shorter overall survival than those with high EWSR1::ETS activity (log-rank test; *P*<0.0001) (**Fig. 1a**). To rule out that the observed difference in overall survival between EWSR1::ET-high and -low is attributable to differences in tumor purity, we repeated the Kaplan-Meier analysis now restricted to samples with an inferred tumor purity of >80%. As shown in **Supplementary Fig. 1a**, patients with low EWSR1::ETS activity retained a significantly lower overall survival probability (*P*=0.0016, log-rank test) even after exclusion of cases with tumor purity ≤80% (n=38). Given the link between low EWSR1::ETS and invasiveness, and as metastatic cells require metabolic adaptation to new niches (Pavlova and Thompson, 2016), we hypothesized that the higher clinical aggressiveness of tumors with low EWSR1::ETS activity could be mediated by metabolic reprogramming. To test this possibility, we carried out a comprehensive time-course metabolomics screen for 159 small metabolites in a EwS cell line (shA-673-1C) harboring a doxycycline (DOX)-induced shRNA against *EWSR1::FLI1* (Tirode *et al*., 2007). As shown in **Fig. 1b**, Kyn – a major endogenous ligand of the AHR receptor (DiNatale *et al*., 2010; Opitz *et al*., 2011) – showed the strongest increase 72 h after *EWSR1::FLI1* knockdown (KD) across159 tested small metabolites (**Supplementary Table 1**). Given these data and as a prior study linked EWSR1::FLI1 with Kyn metabolism and AHR signaling in EwS cell lines (Mutz *et al*., 2016), we focused on Kyn as the top candidate. However, whether the regulatory relationship between the fusion an Kyn and subsequent AHR signaling are operative in EwS patient tumors or in cell line models under physiological conditions (Ceranski *et al*., 2025) remain unclear. Thus, we employed a newly defined pan-tissue AHR activity signature (Sadik *et al*., 2020) comprising 166 genes (see Methods). As shown in **Figs. 1c–e**, consensus K-means clustering of the patients according to their AHR activity scores identified two distinct clusters labeled K1 (AHR-low) and K2 (AHR-high), and demonstrated that patients with high AHR scores (K2) had a significantly shorter overall survival than those with low AHR scores (K1) (log-rank test; *P*=0.031). Patients with high AHR scores (K2) exhibited significantly higher intratumoral expression levels of *AHR* and significantly altered mRNA expression levels of key enzymes (*IL4I1*, *TDO2*, *IDO1*, *KYAT1*) involved in Trp metabolism (**Figs. 1f,g**), indicating a metabolic shift toward increased Trp catabolism and production of the AHR ligand Kyn. In line with this notion, the inferred AHR scores significantly negatively correlated with the previously calculated EWSR1::ETS activity scores (*P*=0.0006, Fisher’s exact test) (**Fig. 1h**). Consistent with these patient data, reanalysis of our previously published Ewing Sarcoma Cell Line Atlas (ESCLA) transcriptome data (Orth *et al*., 2022) revealed a significant upregulation in the AHR signatures across 18 EwS cell lines upon functional EWSR1::ETS KD (**Fig. 1i**). These effects observed in patients and cell lines were mirrored *in vitro* in three-dimensional (3D) spheroids grown in human plasma-like medium (HPLMax) and reduced fetal calf serum (FCS) conditions (7%) (Ceranski *et al*., 2025) in two EwS cell lines harboring DOX-inducible shRNAs against their respective fusions (EWSR1::FLI1 in MHH-ES-1 cells, EWSR1::ERG in TC-106 cells). As shown in **Fig. 1j**, silencing of either fusion for 96 h led to a significant coordinated deregulation of the Trp–Kyn pathway, where the enzymes leading to increased production of Kyn (*TDO2*) or kynurenic acid (*IL4I1*) were upregulated and the and the Kyn-catabolizing enzyme *KMO* was downregulated. Consistent with this, we observed increased Kyn secretion in the supernatants of four EwS cell lines upon silencing of the respective EWSR1::ETS fusion (**Fig. 1k**). Levels of *AHR* itself were not robustly altered by EWSR1::ETS KD (data not shown), which is in line with observations that low *EWSR1::ETS* expression enhances nuclear translocation of AHR rather than its mRNA expression (Mutz *et al*., 2016).

**Figure 1.**
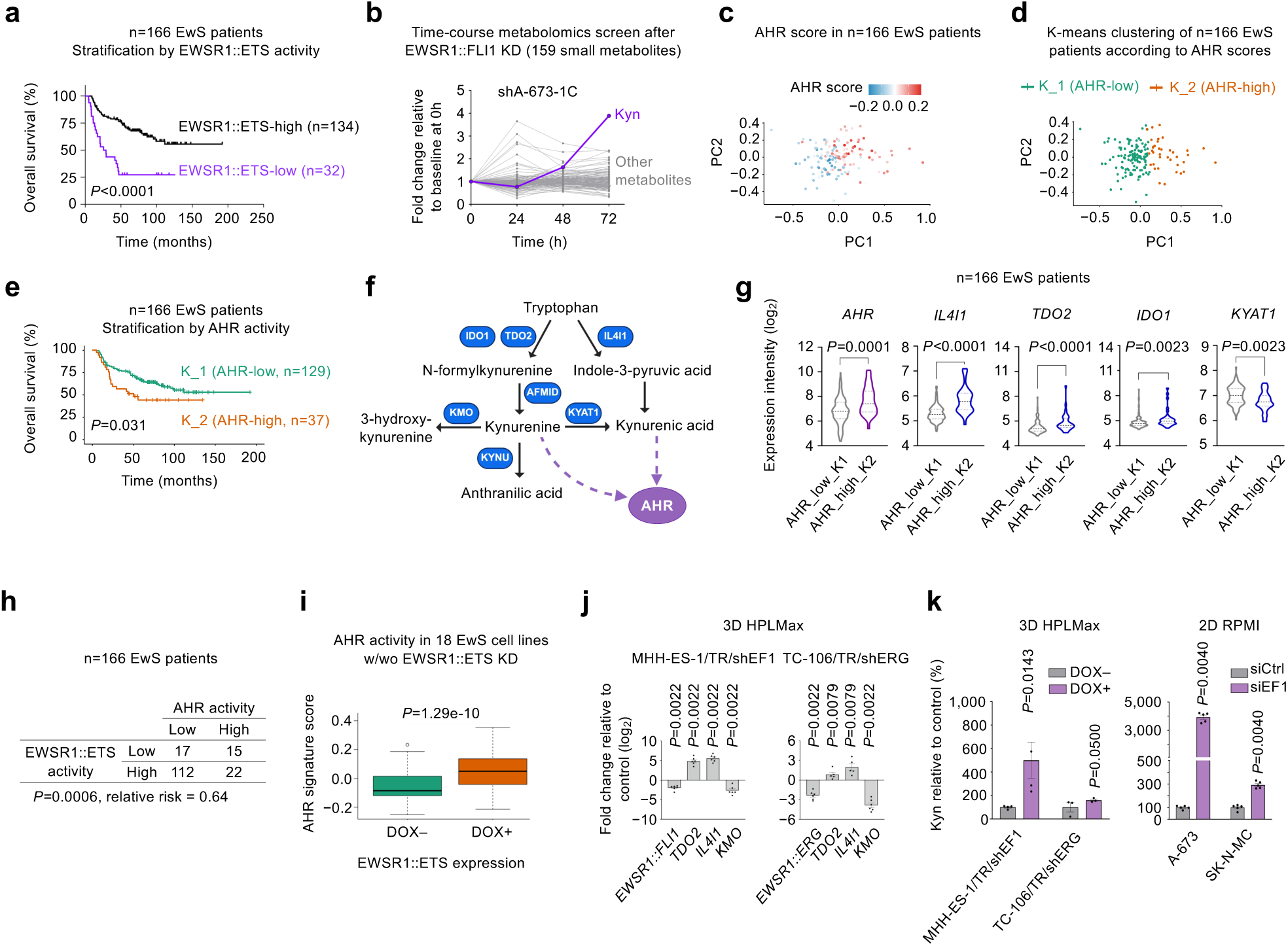
Low EWSR1::ETS activity induces AHR signaling and confers poor overall survival in EwS. **a)** Kaplan-Meier survival analysis of n=166 patients stratified by EWSR1::ETS-activity (log-rank test). **b)** Fold change of 159 small metabolites in a time-course metabolomics screen after *EWSR1::FLI1* KD in shA-673-1C cells. **c,d)** K-means clustering of patients based on AHR activity: K1 (AHR-signaling low) and K2 (AHR-signaling high). **e)** Kaplan-Meier survival analysis of n=166 patients stratified by AHR-activity (low (K1) versus high (K2), log-rank test). **f)** Schematic of the Trp–Kyn–AHR axis. TDO2: tryptophan 2,3-dioxygenase. IL4I1: interleukin-4-induced-1. IDO1: indoleamine 2,3-dioxygenase 1. KYAT1: Kyn aminotransferase 1. KMO: Kyn 3-monooxygenase. KYNU: kynureninase. AHR: aryl hydrocarbon receptor. Created in BioRender. Carreno Gonzalez, M. (2025). https://BioRender.com/kuuj1i8. **g)** Violin plots of the mRNA expression levels of indicated Trp–Kyn–AHR axis factors in 166 EwS tumors stratified by AHR activity. Bold dashed lines represent medians, fine dashed lines quartiles. Two-sided Mann-Whitney test. **h)** Contingency table showing distribution of EWSR1::ETS and AHR activity score-based clusters in EwS tumors (n=166). Two-sided Fisher’s exact test. **i)** Box plots depicting the AHR activity scores of 18 EwS cell lines upon *EWSR1::ETS* KD (DOX+) as compared to controls (DOX–). Horizontal bars represent the median. **j)** Expression levels of *TDO2*, *IL4I1* and *KMO* upon KD of *EWSR1::FLI1* and *EWSR1::ERG* fusions (96 h DOX treatment) in MHH-ES-1/TR/shEF1 and TC-106/TR/shERG, respectively. Horizontal bars represent the mean and whiskers the SEM. n=5−6 biologically independent experiments. Two-sided Mann-Whitney test. **k)** Percentage of Kyn levels relative to control upon fusion KD (left: MHH-ES-1/TR/shEF1 and TC-106/TR/shERG in 3D HPLMax 7% FCS, 96 h DOX treatment; right: A-673 and SK-N-MC in 2D RPMI, 72 h siRNA treatment). Horizontal bars represent the mean and whiskers the SEM. n=3−4 biologically independent experiments. One-sided Mann-Whitney test.

In sum, these data highlight the pronounced impact of EWSR1::ETS on the Trp–Kyn metabolism and extend them to a clinically relevant phenotype in patient tumors with increased AHR signaling and enrichment in EWSR1::ETS-low cells.

### EWSR1::ETS low-induced AHR activity promotes an immunoprotective phenotype in EwS

Since increased AHR activity has previously been linked to an immunosuppressive microenvironment that fostered tumor growth in other cancer types (Quintana *et al*., 2008; Mezrich *et al*., 2010; Liu *et al*., 2018; Neamah, Nagarkatti and Nagarkatti, 2018; Greene *et al*., 2019; Sadik *et al*., 2020), we hypothesized that EwS tumors with increased AHR scores might exhibit an altered composition of immune cell infiltrates. Interestingly, correlating computationally derived Immune Pediatric Signature Scores (IPASS) (Mayoh *et al*., 2023) inferred from our patient transcriptome data with the AHR scores revealed a significantly higher T cell infiltration of AHR-high EwS tumors (**Fig. 2a**). However, a more fine-granular analysis of the immune cell composition (see Methods) revealed not only an enrichment of B cells, (CD8+) T cells, and NK cells (**Fig. 2b**), but also a strong enrichment for immunosuppressive myeloid-derived suppressor cells (MDSCs) and regulatory T cells, whose increased presence is observed in immunotherapy non-responders (Liu *et al*., 2015; Oweida *et al*., 2018; Weber *et al*., 2018; Hou *et al*., 2020; Petrova *et al*., 2023; van Gulijk *et al*., 2023) (**Fig. 2c**). This dual pattern may explain why the outcome of patients with AHR-high scores is worse despite displaying diverse immune infiltration overall. Given that we observed increased Kyn secretion by EWSR1::ETS-low cells and that Kyn can inhibit surface expressed activating receptors (Chiesa *et al*., 2006) and suppress the cytotoxic function of NK cells (Trikha *et al*., 2021), we tested whether EWSR1::ETS levels might influence NK cell activity against EwS cells. To this end, we first carried out NK cell killing assays by co-culturing NK-92 cells with our EwS models with/without DOX-induced silencing of the respective EWSR1::ETS fusion. MHH-ES-1/TR/shEF1 and TC-106/TR/shERG cells were pretreated with or without DOX for 72 h and assessed for cell viability after 24 h of co-culture with NK-92 cells. Excitingly, KD of EWSR1::ETS significantly reduced NK cell-mediated killing of MHH-ES-1/TR/shEF1 and TC-106/TR/shERG EwS cells (**Fig. 2d**). In a second step, although NK-92 cells are widely used as adequate models for NK cell studies (Klingemann, 2023), we repeated these experiments with peripheral blood mononuclear cell (PBMC)-derived primary NK cells from unrelated donors to account for interindividual differences in baseline NK cell activity. Notably, we could fully recapitulate the inhibitory effect of EWSR1::ETS silencing on primary NK cells (**Fig. 2e**). Interestingly, no differential killing effect on EwS cells was detected upon fusion KD when NK cells were removed from the PBMC population (**Fig. 2f**), suggesting that the immunosuppressive effect of low EWSR1::ETS is predominantly mediated through altered NK cells responses.

**Figure 2.**
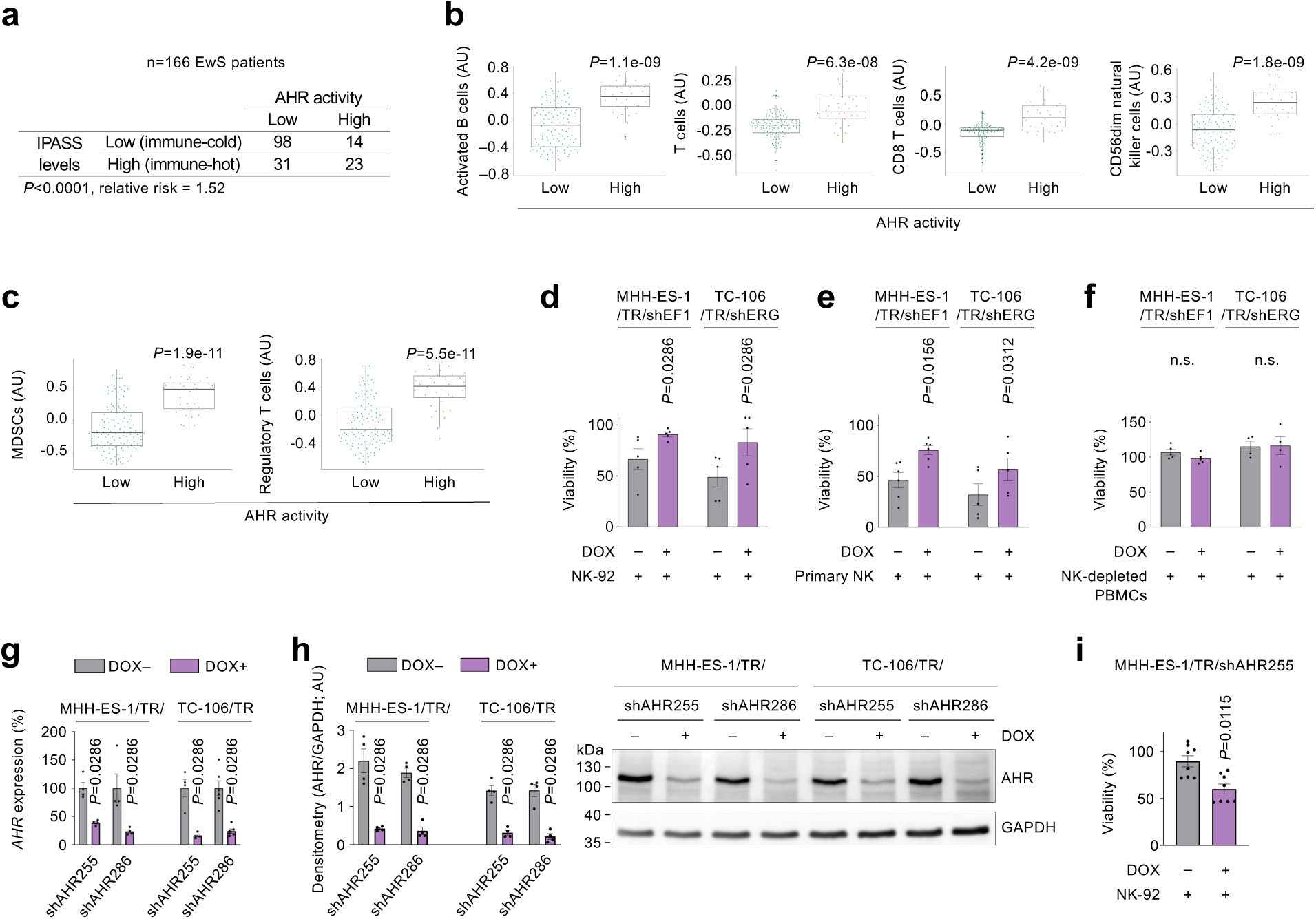
EWSR1::ETS low-induced AHR activity promotes an immunoprotective phenotype in EwS. **a)** Contingency table showing distribution of IPASS score and AHR activity score-based clusters in EwS tumors (n=166). Two-sided Fisher’s exact test. **b,c)** Boxplots showing a comparative analysis of inferred pro-inflammatory immune cell infiltration (b) and immunosuppressive MDSCs and regulatory T cells infiltration (c) of patient tumors in (a) stratified by AHR activity (AHR-low versus -high). Horizontal bars represent the median, boxes the interquartile range, and whiskers the maximum and minimum. Wilcoxon test. **d)** Relative cell viability MHH-ES-1/TR/shEF1 and TC-106/TR/shERG after 24 h co-culture with NK-92 cells. Horizontal bars represent the mean and whiskers the SEM. n=5 independent biological experiments. Two-sided Mann-Whitney test. **e)** Relative cell viability MHH-ES-1/TR/shEF1 and TC-106/TR/shERG after 24 h co-culture with primary NK cells. Horizontal bars represent the mean and whiskers the SEM. n=5–6 independent biological experiments. One-sided paired Wilcoxon test. **f)** Relative cell viability MHH-ES-1/TR/shEF1 and TC-106/TR/shERG after 24 h co-culture with NK cell depleted PBMCs. Horizontal bars represent the mean and whiskers the SEM. n=4–5 independent biological experiments. Two-sided Mann-Whitney test. **g)** Relative *AHR* mRNA (qRT-PCR) and **h**) protein expression (immunoblot) in MHH-ES-1/TR/shAHR and TC-106/TR/shAHR cells upon 96 h of DOX-treatment. AU = arbitrary units. n=4−6 biologically independent experiments. Two-sided Mann-Whitney test. **i)** Relative cell viability MHH-ES-1/TR/shAHR cells after 24 h co-culture with NK-92 cells. Horizontal bars represent the mean and whiskers the SEM, n=6 biologically independent experiments. Two-sided Mann-Whitney test.

To test the possibility that the effect of EWSR1::ETS-low expression on NK cell killing is meditated via AHR signaling in EwS cells, we generated single-cell cloned DOX-inducible shAHR EwS cells, which exhibited a strong and comparable KD of AHR at the mRNA and protein levels across all models (**Figs. 2g,h**). As the experimental setting above, MHH-ES-1/TR/shAHR255 cells were treated with or without DOX for 72 h prior to being seeded at equal numbers and co-cultured with NK-92 cells for additional 24 h. Excitingly, these experiments revealed a strong enhancement of NK cell-mediated killing upon AHR silencing (**Fig. 2i**), which had a comparable magnitude (∼2-fold) to the inhibitory effect of the EWSR1::ETS KD. Collectively, these *in situ* and *in vitro* analyses suggest that AHR activation accounts, at least in part, for the immunoprotective phenotype observed in EWSR1::ETS-low cells.

### AHR promotes proliferation and clonogenicity of EwS cells *in vitro*

Since our results pointed to a role of EWSR1::ETS-mediated metabolic reprogramming of EwS cells concerning Trp–Kyn metabolism in cell-extrinsic phenotypes, we wondered whether this may also have cell-intrinsic effects. To test this possibility, we subjected our single cell-cloned shAHR EwS models to a series of *in vitro* assays in physiological medium (HPLMax) (Cantor *et al*., 2017; Vande Voorde *et al*., 2019; Ceranski *et al*., 2025). As shown in **Fig. 3a**, conditional AHR silencing significantly reduced proliferation of MHH-ES-1 and TC-106 cells in short-term assays (96 h), which was accompanied by delayed cell cycle progression at G1 (**Fig. 3b**). To assess the cell-intrinsic effects of AHR for a longer period of time (10–12 d), we carried out colony-formation assays. Strikingly, KD of AHR almost completely suppressed clonogenic growth of both EwS cell lines (**Fig. 3c**), an effect not observed for control cells expressing a DOX-inducible non-targeting shRNA (**Supplementary Fig. 1b**). Given these strong effects and to exclude a potential bias through clonal artefacts, we repeated key assays with the parental bulk-level shAHR cells, which exhibited similar remaining AHR levels upon DOX-treatment, and comparable phenotypes regarding cell proliferation and clonogenicity as well as spheroidal growth (**Fig. 3d**, **Supplementary Figs. 1b–d**). To examine whether similar effects could be achieved via pharmacological inhibition and to explore the potential of AHR as a drug target in EwS, we subjected MHH-ES-1 and TC-106 cells to ascending doses of the specific small molecule AHR antagonist GNF-351 (Smith *et al*., 2011; Van Den Bogaard *et al*., 2015) and carried out short-term viability and long-term colony formation assays, which recapitulated our AHR KD results (**Figs. 3e,f**). Taken together, these results point to an important role of AHR as a druggable target promoting proliferation of EwS cells *in vitro*.

**Figure 3.**
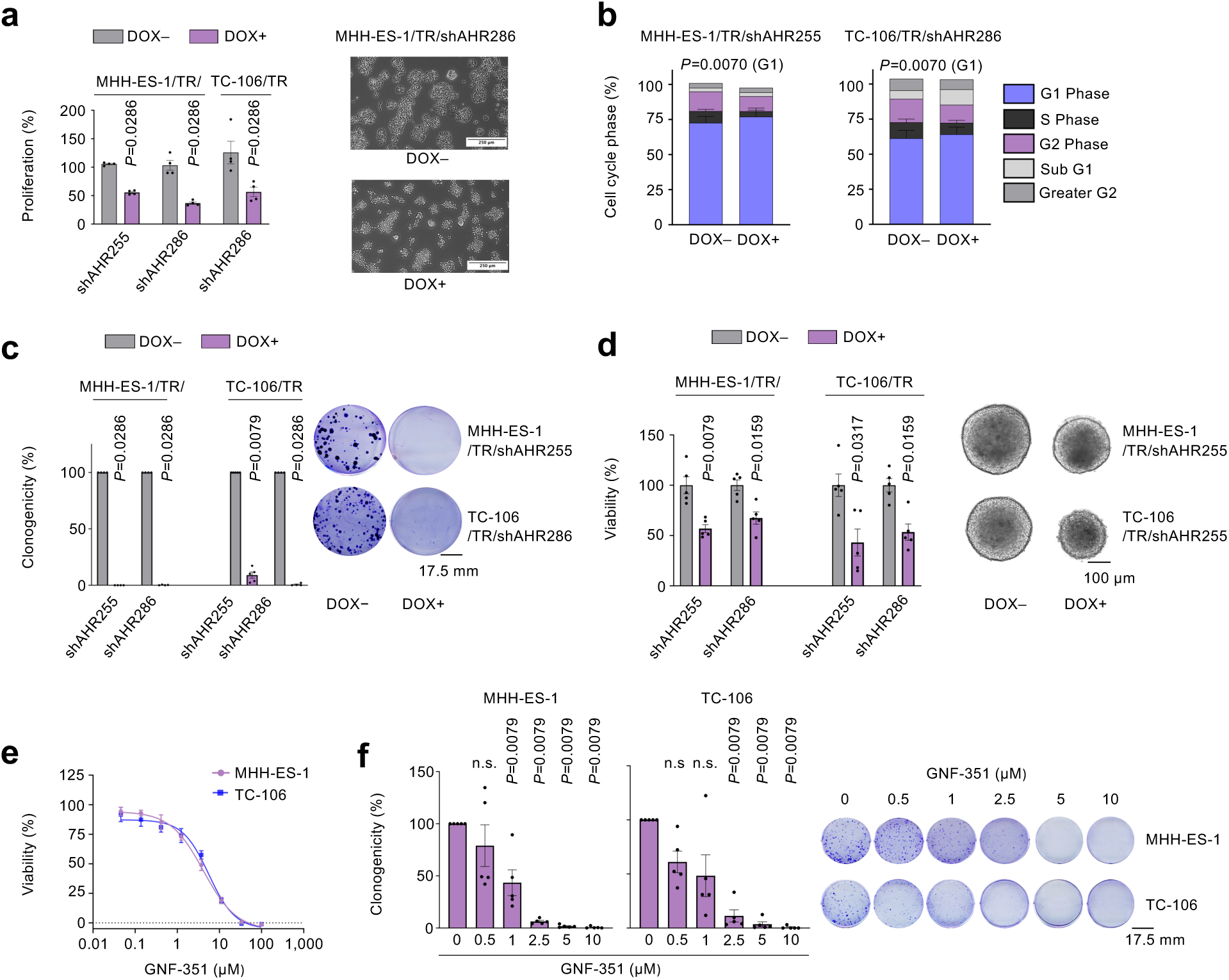
AHR promotes proliferation and clonogenicity of EwS cells *in vitro*. **a)** Proliferation of MHH-ES-1/TR/shAHR cells following 96 h of DOX-induced AHR KD (left) and representative images of cell proliferation of clone MHH-ES-1/TR/shAHR (5× objective, right). Horizontal bars represent the mean and whiskers the SEM. n=4 biological independent experiments. Two-sided Mann-Whitney test. **b)** Analysis of the cell cycle by PI staining at 96 h post-DOX treatment in MHH-ES-1/TR/shAHR255 cells and TC-106/TR/shAHR286. n=5 biologically independent experiments. Two-sided Mann-Whitney test (G1 phase). **c)** Relative clonogenicity of single clones MHH-ES-1/TR/shAHR255 and shAHR286 and TC-106/TR/shAHR255 and shAHR286 cells following 10–12 d of DOX-treatment (left) and representative colony images (right). Horizontal bars represent the mean and whiskers the SEM. n=4−5 biologically independent experiments. Two-sided Mann-Whitney test. **d)** Relative clonogenicity of MHH-ES-1/TR/shAHR and TC-106/TR/shAHR cells containing a DOX-inducible KD construct for targeting AHR (sh255 and 286) and non-targeting control (shControl) for 10–12 d (left) and representative colony images (right). Horizontal bars represent the mean and whiskers the SEM, n=4 biologically independent experiments. Two-sided Mann-Whitney test. **e)** Dose-response curves of MHH-ES-1 and TC-106 cells treated with GNF-351 for 72 h. Relative viability was determined by resazurin assay. n=5 biologically independent experiments. **f)** Relative clonogenicity of MHH-ES-1 and TC-106 cells following 72 h GNF-351 treatment (left) and representative colony images (right) as compared to DMSO controls. n=5 biologically independent experiments. Horizontal bars represent the mean and whiskers the SEM. Two-sided Mann-Whitney test.

### AHR promotes tumor growth and metastasis of EwS *in vivo*

Given the strong phenotypes observed upon AHR silencing in our *in vitro* models grown in physiological conditions, we next tested the impact of AHR on tumor growth and metastasis *in vivo*. To that end, we inoculated our single cell-cloned shAHR EwS cells subcutaneously into the right flank of NOD/Scid/gamma (NSG) mice and started DOX-treatment once tumors were palpable (**Fig. 4a**). As shown in **Fig. 4b**, conditional KD of AHR led to tumor regression in both MHH-ES-1- and TC-106-derived xenografts. In a next step, we employed a spontaneously metastasizing orthotopic model by inoculating MHH-ES-1/TR/shAHR cells into the tibial plateau of NSG mice and started DOX-treatment the subsequent day (Cidre-Aranaz *et al*., 2022) (**Fig. 4c**). Mice were killed at individual days once predefined humane endpoints, such as signs of tumor-associated limping of the injected leg or abdominal distension as a possible sign of metastasis, were detectable. At the endpoint, there was a significant reduction in the weight of the primary tumors of the *AHR*-silenced DOX+ group, as determined by surrogate measure of the weight of the inoculated leg (**Fig. 4d**). Further, necropsy of the animals showed that there was a significant difference in liver weights concordant with the significantly reduced number of macroscopically detectable liver metastases in the AHR KD group (**Figs. 4e,f**). Collectively, these *in vivo* results further emphasize the important role of AHR signaling for tumor progression of EwS.

**Figure 4.**
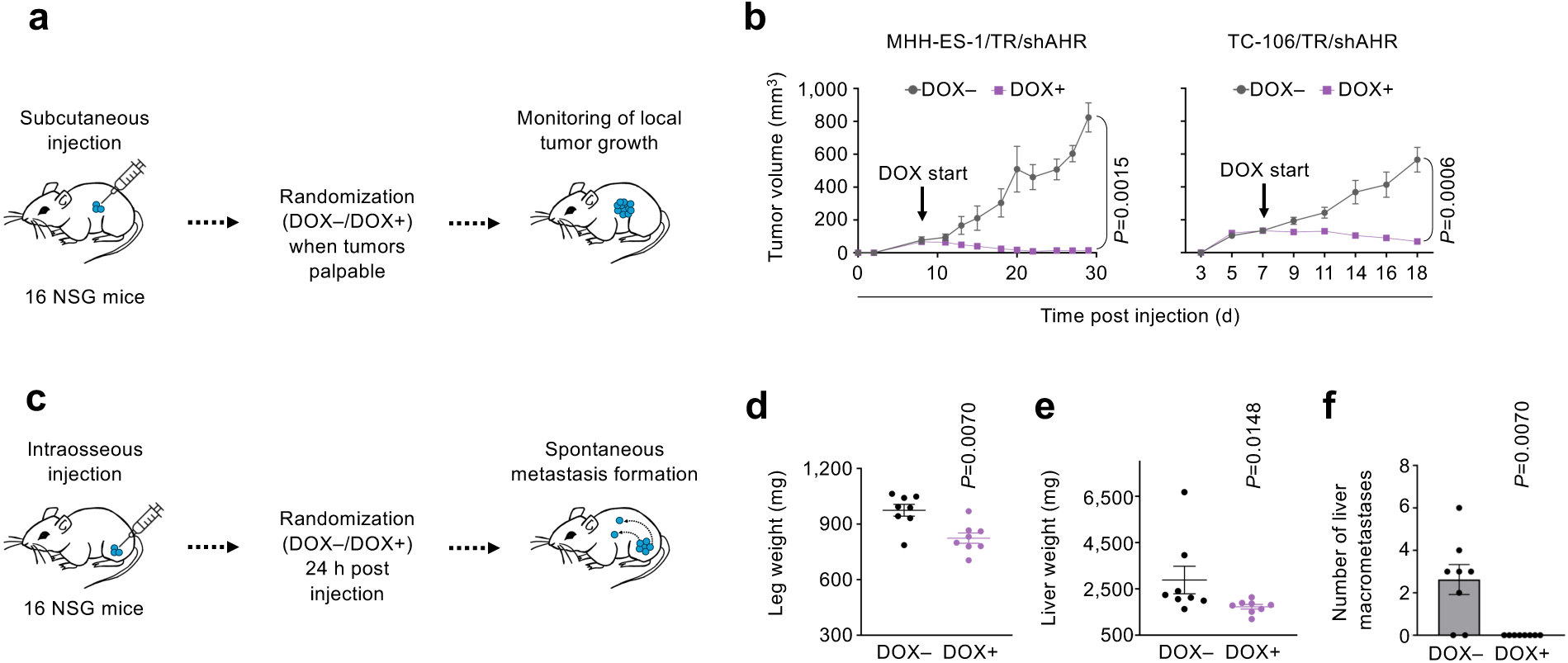
AHR promotes tumor growth and metastasis of EwS *in vivo*. **a)** Schematic of subcutaneous (s.c.) injection experiments in mice. Mice were randomized once tumors were palpable (day 5 for MHH-ES-1 cells; day 7 for TC-106 cells) into control (DOX–) and AHR KD groups (DOX+). **b)** Analysis of volumes of subcutaneous tumors over time. Dots represent the mean and whiskers the SEM. n=8 biologically independent animals per group. Multigroup unpaired two-sided Mann-Whitney test. **c)** Schematic of orthotopic injection experiments. MHH-ES-1/TR/shAHR cells were injected in the tibial plateau of NSG mice that were randomized 1 d thereafter in the control group (DOX–) and AHR KD group (DOX+). **d)** Analysis of leg weight (mg) of both groups as depicted in (c) at the day of endpoint. Horizontal line indicates the median and whiskers the SEM. n=8 animals/group. Two-sided Mann-Whitney test. **e)** Liver weight (mg) of both groups as depicted in (c) at the day of the endpoint. Horizontal line indicates the median and whiskers the SEM. n=8 animals/group. Two-sided Mann-Whitney test. **f)** Number of liver macrometastases of both groups as depicted in (c). Horizontal bars represent the mean and whiskers the SEM. n=8 animals/group. Two-sided Mann-Whitney test.

## DISCUSSION

EwS remains one of the most aggressive pediatric cancers, characterized by early metastatic spread and high relapse rates (Zöllner *et al*., 2021). Unlike other difficult-to-treat malignancies, EwS has thus far responded poorly to immunotherapy (Tawbi *et al*., 2017) largely due to its immunologically ‘cold’ tumor microenvironment (Cillo *et al*., 2022; Kuo *et al*., 2025). Tumor metabolism has emerged as a key regulator of immune surveillance, yet the extent to which the EwS-defining EWSR1::ETS fusions shape metabolic–immune crosstalk has been unclear.

Our study identifies the Trp–Kyn–AHR axis as a central mediator of EwS progression and immune evasion. While aberrant Trp catabolism and AHR activation have been implicated in immune suppression across diverse cancers (Opitz *et al*., 2011; Ye *et al*., 2018; Solvay *et al*., 2023; Holfelder *et al*., 2024) and AHR nuclear translocation upon silencing of EWSR1::FLI1 was reported in EwS cell lines (Mutz *et al*., 2016), we show for the first time that reduced EWSR1::ETS fusion activity and consequently elevated AHR activity in EwS tumors is associated with inferior patient overall survival and a complex immune phenotype marked by signatures of both activated lymphocytes and immunosuppressive cell populations. This duality suggests that AHR activation may enable EwS tumors to sustain immune infiltration while simultaneously establishing resistance mechanisms.

Functionally, we found that low *EWSR1::ETS* expression promoted NK cell resistance pointing to an immune evasion state whose precise underpinnings remain to be defined. While it has been established that Kyn has a direct negative impact on NK cell function (Chiesa *et al*., 2006; Trikha *et al*., 2021), it will require further dissection whether this reflects impaired NK cell cytotoxicity, intrinsic tumor resistance, or both. Prior work has highlighted several EwS immune-evasion strategies—including high CD47 expression, altered calreticulin dynamics (Luo *et al*., 2024), and upregulation of non-classical HLA molecules (Spurny *et al*., 2017; Altvater *et al*., 2021)—but their regulation by fluctuations in EWSR1::ETS fusion levels has not been explored.

Beyond its immune effects, we demonstrate that AHR activity directly sustains EwS tumor growth. AHR silencing impaired proliferation, clonogenicity, cell cycle progression and spheroidal growth in vitro, and in xenograft and orthotopic models significantly reduced tumor burden and metastasis. These findings align with evidence from other cancers linking AHR signaling to proliferation, survival, and invasiveness (Puga, Xia and Elferink, 2002; Peng *et al*., 2009; Murray, Patterson and Perdew, 2014; Stanford *et al*., 2016). While these pro-tumorigenic effects of AHR have been linked to reduced cell-to-cell contact by downregulation of E-cadherin and increased cell motility and survival (Dietrich and Kaina, 2010; Opitz *et al*., 2011; D’Amato *et al*., 2015; Novikov *et al*., 2016; Shadboorestan *et al*., 2019), further work is needed to pinpoint the precise pro-tumorigenic mechanisms in EwS, including potential roles for altered cell adhesion, epithelial-to-mesenchymal–like programs, and survival signaling pathways. The implementation of humanized mice and/or mice with functional NK cells will be essential for clarifying how AHR activity integrates tumor-intrinsic effects with the immune compartment.

Therapeutically, our findings position the Trp–Kyn–AHR axis as an attractive druggable target in EwS. Upstream enzymes (IDO1, TDO2, IL4I1) appear to contribute to AHR activation in EWSR1::ETS-low cells, although targeting these individually has proven challenging: While several IDO1 inhibitors such as epacadostat and navoximod have been investigated in combination with PD-1/PD-L1 targeting therapies, these failed to demonstrate clinical benefit in trials for metastatic melanoma (Long *et al*., 2019) or advanced solid tumors (Jung *et al*., 2019), likely due to structural resemblance of the IDO1 inhibitor to AHR agonists (Lewis *et al*., 2017; Moyer *et al*., 2017; Long *et al*., 2019) and/or compensation by IDO1 inhibitors such as IL4I1 (Sadik *et al*., 2020). Few selective and potent inhibitors, however, have been described for TDO2, none of which have progressed to clinical trial (Kozlova and and Frédérick, 2019) and some additionally act as AHR agonists and are therefore of limited use in this context (Opitz *et al*., 2020). The link between activation of AHR by metabolites produced downstream of IL4I1 and tumor cell immune evasion has only recently been recognized (Sadik *et al*., 2020), and inhibitors are therefore still in an early stage of development (MacKinnon *et al*., 2020; Hess *et al*., 2025).

By contrast, direct AHR inhibition may circumvent these limitations. Indeed, the AHR antagonist GNF-351 mirrored the effects of genetic silencing, impairing EwS proliferation and survival. The development of next-generation selective AHR antagonists or tunable modulators could enable effective targeting with reduced systemic toxicity, warranting systematic in vivo testing. Indeed, efforts are underway to develop selective AHR inhibitors, including AHR modulating compounds that fine-tune rather than fully antagonize AHR activity to limit potential systemic effects (Yin *et al*., 2012; Kober *et al*., 2023; Aggen *et al*., 2024).

In conclusion, we uncover a previously unrecognized role for the EWSR1::ETS-regulated Trp– Kyn–AHR axis in EwS progression and immune evasion. AHR emerges not only as a critical regulator of EwS metabolism and cell-intrinsic growth but also as a mediator of resistance to NK cell cytotoxicity. By linking oncogenic transcriptional control to metabolic rewiring and immune evasion, our study highlights AHR as an actionable therapeutic node with potential to improve outcomes for patients with metastatic and relapsed EwS.

## CONFLICT OF INTEREST

Authors of this manuscript have patents on AHR inhibitors in cancer (WO2013034685, C.A.O.); A method to multiplex tryptophan and its metabolites (WO2017072368, C.A.O.); A transcriptional signature to determine AHR activity (WO2020208190, A.S., C.A.O.); Interleukin-4-induced gene (IL4I1) and its metabolites as biomarkers for cancer (WO2021116357, A.S., C.A.O.). The other authors declare no conflict of interest.

## AUTHOR CONTRIBUTIONS

M.J.C.G. carried out the majority of the *in vitro* and *in vivo* experiments. K.M.H, A.K.C., F.H.G., Z.A.K., M.Z. and A.R. helped in molecular cloning and *in vitro* assays. S.O. and M.M.L.K. helped in NK-cell experiments and R.I., A.B., and F.C.A. in *in vivo* experiments. A.S., C.H., and A.C.E. carried out bioinformatics analyses. A.H. and M.J.C.G. carried out measurements of Kyn levels. A.J., D.S., and O.D. provided metabolomics data. M.M.L.K. and C.A.O. provided experimental guidance. M.J.C.G., K.M.H. and T.G.P.G. wrote the paper and designed the figures. T.G.P.G. conceived and supervised the study and provided laboratory infrastructure. All authors read and approved the final manuscript.

## ACKNOWLEDGEMENTS

We thank S. Kutschmann, F. Zahnow, N. Gmelin, and S. Knoth for excellent technical assistance. We also thank Prof. Dr. A. Cerwenka and Dr. A. Stojanovic (Department of Immunobiochemistry, Mannheim Institute for Innate Immunoscience, Medizinische Fakultät Mannheim der Universität Heidelberg, Germany) for consultations regarding NK cell experiments. We thank members of the Genetics and Biology of Pediatric Cancer laboratory and of Inserm U830 for fruitful discussion. and we are grateful to Metabolon Inc. for their expert assistance and support in conducting the metabolomics experiments.

## FUNDING

This work was mainly supported by grants from the Boehringer-Ingelheim foundation, the German Cancer Consortium (DKTK, PredictAHR), and the German Cancer Aid (DKH-70115315). The laboratory of T.G.P.G. is supported by grants from the Matthias-Lackas foundation, the Dr. Leopold und Carmen Ellinger foundation, Dr. Rolf M. Schwiete foundation (2021-007; 2022-031), the German Cancer Aid (DKH-70114278, DKH-70115914), the SMARCB1 association, the Ministry of Education and Research (BMBF; SMART-CARE and HEROES-AYA), the KiKa foundation, the Fight Kids Cancer foundation (FKC-NEWtargets), the KiTZ-Foundation in memory of Kirstin Diehl, the KiTZ-PMC twinning program, Sarcoma Research UK, and the Barbara and Wilfried Mohr foundation. The laboratory of T.G.P.G. is co-funded by the European Union (ERC, CANCER-HARAKIRI, 101122595). Views and opinions expressed are however those of the authors only and do not necessarily reflect those of the European Union or the European Research Council (ERC). Neither the European Union nor the granting authority can be held responsible for them. A.R., F.H.G., and A.C.E. were supported by scholarships of the German Cancer Aid and the German Academic Scholarship Foundation. M.Z. also acknowledges funding by the German Academic Scholarship Foundation as well as by the Kind-Philipp foundation. K.M.H. was supported by a fellowship of the Alexander von Humboldt foundation. C.A.O. acknowledges support from the ERC under the European Union’s Horizon 2020 research and innovation programme (grant agreement number 101045257, CancAHR). Further, this work in the laboratory of O.D. was supported by grants from Ligue Nationale Contre le Cancer (Equipe labellisée). A.J. was supported by a PhD fellowship from Institut Curie.

## METHODS

### Cell lines and cell culture conditions

Human HEK293T, MHH-ES-1, and NK-92 cell lines were provided by the German collection of Microorganisms and Cell cultures (DSMZ). The human EwS cell line TC-106 was kindly provided by the Children’s Oncology Group (COG). The standard cell culture conditions for HEK293T cells and EwS cells for viral transduction were as follows: RPMI 1640 with stable glutamine (ThermoFisher Scientific, USA), 10% tested doxycycline (DOX)-free FCS, penicillin (100 U/mL), and streptomycin (100 μg/mL) (1% penicillin/streptomycin) (Sigma-Aldrich, Germany). Newly established physiological conditions that better recapitulate the *in situ* situation were used for performing all other EwS experiments using HPLM (ThermoFisher Scientific, USA) plus unique Plasmax components, 7% FCS, and 1% penicillin/streptomycin (Ceranski *et al*., 2025).

To culture the NK-92 cell line, the base medium was composed of RPMI 1640 medium with 2 mM L-glutamine, heat-inactivated FCS (56 °C for 30 min), and 1% penicillin/streptomycin. Lyophilized IL-2 was resuspended in 50 μL 100 mM acetic acid and 50 μL of 0.2% FCS in PBS (10^6^ U/mL) and stored at −80 °C until use. The daily cell culture media was prepared by combining 50 mL of the base medium with 500 μL of HEPES 1 M (final concentration 10 mM) and 5 μL IL-2 (final concentration 10 ng/mL). NK-92 cells were maintained at confluency of 1−4×10^5^ cells/mL. All cell lines were incubated at 37 °C and 5% CO_2_ in a controlled humidified environment.

All cell cultures were regularly tested for mycoplasma contamination (Mycoplasma PCR Detection Kit, Applied Biological Materials) and identified by SNP (single nucleotide polymorphism) or STR (short tandem repeat)-profiling (Multiplexion, Heidelberg, Germany).

### Metabolomics analysis

For the metabolic analysis, shA-673-1C cells that enable a DOX-inducible expression of a specific shRNA against *EWSR1::FLI1* (Tirode *et al*., 2007) were cultured in the presence or absence of DOX. Cells were plated on day -7, and DOX was added at 72 h before and from then on every 12 h until harvest,. These time points were selected to capture early and intermediate effects of EWSR1::FLI1 inhibition. Cells were harvested as follows: Medium was removed, monolayers were rinsed with PBS and cells were detached with trypsin for 5 min at 37 °C. Six volumes of medium were added to the trypsin–cell mixture, and cells were resuspended by gentle pipetting. Cells were counted, and aliquots were taken for RNA isolation (3.3% of total cells) and protein lysate preparation (6.6%). The remaining cells were pelleted at 600g for 3 min in polystyrene tubes, washed once with PBS, snap-frozen in liquid nitrogen, and stored at −80 °C until analysis.

Metabolomic profiling was performed in collaboration with Metabolon (Durham, NC, USA) using both liquid chromatography–mass spectrometry (LC–MS) and gas chromatography–mass spectrometry (GC–MS). The following steps were carried out at Metabolon:

#### Sample Preparation

The sample preparation process was carried out using the automated MicroLab STAR® system from Hamilton Company. Recovery standards were added prior to the first step in the extraction process for QC purposes. Sample preparation was conducted using a proprietary series of organic and aqueous extractions to remove the protein fraction while allowing maximum recovery of small molecules. The resulting extract was divided into two fractions; one for analysis by LC and one for analysis by GC. Samples were placed briefly on a TurboVap® (Zymark) to remove the organic solvent. Each sample was then frozen and dried under vacuum. Samples were then prepared for the appropriate instrument, either LC/MS or GC/MS.

#### Liquid chromatography/Mass Spectrometry (LC/MS, LC/MS2)

The LC/MS portion of the platform was based on a Waters ACQUITY UPLC and a Thermo-Finnigan LTQ mass spectrometer, which consisted of an electrospray ionization (ESI) source and linear ion-trap (LIT) mass analyzer. The sample extract was split into two aliquots, dried, then reconstituted in acidic or basic LC-compatible solvents, each of which contained 11 or more injection standards at fixed concentrations. One aliquot was analyzed using acidic positive ion optimized conditions and the other using basic negative ion optimized conditions in two independent injections using separate dedicated columns. Extracts reconstituted in acidic conditions were gradient eluted using water and methanol both containing 0.1% formic acid, while the basic extracts, which also used water/methanol, contained 6.5 mM ammonium bicarbonate. The MS analysis alternated between MS and data-dependent MS2 scans using dynamic exclusion.

#### Gas chromatography/mass spectrometry (GC/MS)

The samples destined for GC/MS analysis were re-dried under vacuum desiccation for a minimum of 24 h prior to being derivatized under dried nitrogen using bistrimethyl-silyl-triflouroacetamide (BSTFA). The GC column was 5% phenyl and the temperature ramp is from 40° to 300° C in a 16 min period. Samples were analyzed on a Thermo-Finnigan Trace DSQ fast-scanning single-quadrupole mass spectrometer using electron impact ionization. The instrument was tuned and calibrated for mass resolution and mass accuracy on a daily basis. The information output from the raw data files was automatically extracted as discussed below.

#### Accurate mass determination and MS/MS fragmentation (LC/MS/MS)

The LC/MS portion of the platform was based on a Waters ACQUITY UPLC and a Thermo-Finnigan LTQ-FT mass spectrometer, which had a linear ion-trap (LIT) front end and a Fourier transform ion cyclotron resonance (FT-ICR) mass spectrometer backend. For ions with counts greater than 2 million, an accurate mass measurement could be performed. Accurate mass measurements could be made on the parent ion as well as fragments. The typical mass error was less than 5 ppm. Ions with less than two million counts require a greater amount of effort to characterize. Fragmentation spectra (MS/MS) were typically generated in data dependent manner, but if necessary, targeted MS/MS could be employed, such as in the case of lower level signals.

The informatics system consisted of four major components, the Laboratory Information Management System (LIMS), the data extraction and peak-identification software, data processing tools for QC and compound identification, and a collection of information interpretation and visualization tools for use by data analysts. The hardware and software foundations for these informatics components were the LAN backbone, and a database server running Oracle 10.2.0.1 Enterprise Edition.

#### Data Extraction and Quality Assurance

The data extraction of the raw mass spec data files yielded information that could loaded into a relational database and manipulated without resorting to BLOB manipulation. Once in the database the information was examined, and appropriate QC limits were imposed. Peaks were identified using Metabolon’s proprietary peak integration software, and component parts were stored in a separate and specifically designed complex data structure.

#### Compound identification

Compounds were identified by comparison to library entries of purified standards or recurrent unknown entities. Identification of known chemical entities was based on comparison to metabolomic library entries of purified standards. As of this writing, more than 1,000 commercially available purified standard compounds had been acquired registered into LIMS for distribution to both the LC and GC platforms for determination of their analytical characteristics. The combination of chromatographic properties and mass spectra gave an indication of a match to the specific compound or an isobaric entity. Additional entities could be identified by virtue of their recurrent nature (both chromatographic and mass spectral). These compounds have the potential to be identified by future acquisition of a matching purified standard or by classical structural analysis.

#### Curation

A variety of curation procedures were carried out to ensure that a high-quality data set was made available for statistical analysis and data interpretation. The QC and curation processes were designed to ensure accurate and consistent identification of true chemical entities, and to remove those representing system artifacts, mis-assignments, and background noise. Metabolon data analysts use proprietary visualization and interpretation software to confirm the consistency of peak identification among the various samples. Library matches for each compound were checked for each sample and corrected if necessary.

#### Normalization

For studies spanning multiple days, a data normalization step was performed to correct variation resulting from instrument inter-day tuning differences. Essentially, each compound was corrected in run-day blocks by registering the medians to equal one and normalizing each data point proportionately (termed the ‘block correction’).

### AHR signature score in ESCLA EwS cell lines

#### Gene Expression Data

Expression data for 18 EwS cell lines were obtained from the Ewing Sarcoma Cell Line Atlas (ESCLA) (Orth *et al*., 2022). Samples were profiled on Affymetrix Clariom D arrays and processed using robust multi-array average (RMA) normalization. Log₂-transformed signal intensities were used for downstream analyses.

#### AHR Signature Score

GSVA (v2.2.0) was applied to compute enrichment scores for the pan-tissue AHR transcriptional signature defined by Sadik *et al*. (Sadik *et al*., 2020). One score per sample was derived, reflecting AHR pathway activity.

#### Statistical Analysis

Differential AHR activity between EWSR1::ETS KD and control (non-KD) conditions was tested using linear modeling with the limma package (3.64.3), including cell line as a blocking factor. Results were visualized as boxplots.

### Survival analysis of EwS patient cohort and AHR signaling and specific type of immune cell infiltration analysis

The microarray data of 166 EwS tumors of the following datasets GSE63157, GSE34620, GSE12102, GSE17618 (Scotlandi *et al*., 2009; Savola *et al*., 2011; Postel-Vinay *et al*., 2012; Volchenboum *et al*., 2015) were downloaded from the Gene Expression Omnibus database (https://www.ncbi.nlm.nih.gov/geo/). The datasets generated on the Affymetrix Human Exon 1.0 ST Array were preprocessed and normalized using the *AffyGEx* package (https://github.com/ahmedasadik/AffyGEx). Datasets generated on the Affymetrix Human Genome U133 Plus 2.0 Array were processed and normalized using *affy* (Gautier *et al*., 2004) and *gcrma* (Irizarry *et al*., 2003). The normalized expression matrices were z-transformed and merged into a single expression matrix and batch corrected using principle component analysis and the *removeBatchEffect* function from *limma* (Ritchie *et al*., 2015). The batch corrected matrix was used to calculate a single sample AHR activity score as described previously (Sadik *et al*., 2020), in addition to performing weighted gene co-expression network analysis (WGCNA) (Langfelder and Horvath, 2008). To identify WGCNA modules that associate with AHR activity, we correlated the WGCNA modules with the AHR activity scores, and performed a global test using the WGCNA modules as predictors of the AHR activity score. Module selection was set at a Pearson correlation coefficient threshold of ± 0.2 at a *P*-value of 0.05 (Evans, 1996), and at a *P*-value of 0.05 for the global test (AHR associated modules, AAMs). K-means consensus clustering (Wilkerson and Hayes, 2010) was performed using the AAMs to identify the number of clusters. The Kaplan-Meier method and log-rank test were used to evaluate the association of the EwS AHR clusters with survival outcome (Therneau and Grambsch, 2000; Therneau, 2024). Single sample immune infiltration scores were generated using MCP-counter, and by applying GSVA (Hänzelmann, Castelo and Guinney, 2013) to the immune signatures (Charoentong *et al*., 2017). The Wilcoxon rank sum test was used for all pairwise group comparisons.

### Single-sample gene set enrichment analysis (ssGSEA)

To quantify sample-based transcriptomic signature enrichment in the EwS patient cohort (n=166) ssGSEA was employed (Barbie *et al*., 2009; Hänzelmann, Castelo and Guinney, 2013). EWSR1::ETS fusion activity signature was assessed using the IC-EwS gene signature previously established (Aynaud *et al*., 2020). The resulting ssGSEA scores were used to classify tumor samples into EWSR1::ETS-high and EWSR1::ETS-low groups via K-means clustering. The *tidyestimate* package (v1.1.1) in R was used to infer tumor purity. Stromal, immune, ESTIMATE, and tumor purity scores were derived from the normalized gene expression data of the n=166 EwS patient tumors using the ESTIMATE algorithm (Yoshihara *et al*., 2013). Similarly, T-cell infiltration levels were inferred using the Immune Pediatric Signature Score (IPASS), a validated 15-gene immune signature associated with CD8+ T-cell infiltration in high-risk pediatric tumors (Mayoh *et al*., 2023). Resulting immune infiltration scores were subsequently used to classify tumors into T cell infiltration-high and T cell infiltration-low groups. To examine potential relationships between AHR pathway activation, EWSR1::ETS fusion activity, and immune cell infiltration 2×2 contingency table with two-sided Fisher’s exact tests were employed.

### Spheroidal growth for 3D assays

EwS spheres were grown on high gel-strength agar coated T-25 flasks as previously described (Ceranski *et al*., 2025). For MHH-ES-1/TR/shEF1 1.8×10^6^ cells per T-25 were seeded and for TC-106/TR/shERG, 8×10^5^ cells per T-25. Spheroids were treated with 1 μg/mL DOX for 96 h, with DOX refreshed every 48 h, then spheroids collected for RNA isolation or the supernatant collected for Kyn level measurement.

### RNA isolation, reverse transcription, and qRT-PCR

To determine KD efficiency and gene expression after treating cells with DOX, RNA isolation was performed using the NucleoSpin® RNA kit as per manufacturer’s instructions (Macherey-Nagel, Germany), followed by cDNA synthesis using 1,000 ng/μL or 500 ng/μL RNA with the High-Capacity cDNA Reverse Transcription Kit (Applied Biosystems, USA) as per manufacturer’s instructions. qRT-PCR reactions were performed using SYBR Select master mix (Applied Biosystems, USA) with 1:5 diluted cDNA and 0.5 μM forward and reverse primers adjusted to a reaction with total volume of 15 μL. Reactions were performed on a CFX Opus 96 (Bio-Rad, USA) and analyzed using Bio-Rad CFX Maestro 2.3 version software. Gene expression was normalized to the *RPLP0* housekeeping gene and fold change calculated by the 2^ΔΔ^CT method (Livak and Schmittgen, 2001). Oligonucleotides and primers were purchased from Eurofins genomics (Sigma-Aldrich) and are listed in **Supplementary Table 2**. The thermal conditions for qRT-PCR were as follows: heat activation at 95 °C for 2 min (1 cycle), DNA denaturation at 95 °C for 10 sec, and annealing and extension at 60 °C for 10 sec (total 50 cycles). Final denaturation at 95 °C for 30 sec, annealing at 65 °C for 30 sec, and extension at 65−95 °C (increase 0.5 °C/5 sec) 1 cycle.

### Ultra-High-Performance LC to measure Kyn

Kyn levels were measured in the supernatant of MHH-ES-1/TR/shEF1 or TC-106/TR/shERG EwS cells in EWSR1::ETS-high and -low context, in the established physiological conditions (3D, HPLMax, 7% FCS, 1% penicillin/streptomycin). DOX treatment was performed for 96 h, and refreshed every 48 h. Cells were grown in spheres as described before. In summary, spheres and supernatant were collected in 15 mL reaction tubes and centrifuged for 4 min, 1,200 rpm, room temperature. A record of the total volume of each sample was taken for normalization. 200 μL of supernatant were collected in reaction tubes and snap frozen in liquid nitrogen immediately. The rest of the supernatant was discarded, and spheres were further collected for protein quantification. Spheres were lysed in RIPA buffer (Serva electrophoresis, Germany) for 10 min and protein was quantified with Pierce BCA Protein Assay Kit (ThermoFisher Scientific, USA). Supernatant was stored in –80 °C until further analysis. For the sample preparation for Kyn measurement, 33.73 μL of trichloroacetic acid was added to 200 μL of each sample, vortexed to mix, and incubated for 15 min. After, samples were centrifuged at 4 °C, 15,000 g for 10 min and supernatant transferred to 250 μL conical point glass inserts for UHPLC (Agilent Technologies, USA). Measurements were performed in an UltiMate 3000 UHPLC (ThermoFisher Scientific). Metabolites were separated by reversed-phase chromatography on an Accucore™ aQ C18 column (ThermoFisher Scientific) 2.6 µm, 2.1 × 150 mm heated to 35 °C and equilibrated with five column volumes of 98% solvent A (H_2_O with 0.1% trifluoroacetic acid) with a flow rate of 0.55 mL/min. Separation of Trp and Kyn was achieved by increasing concentration of solvent B (acetonitrile with 0.1% trifluoroacetic acid) in solvent A with the following gradient: 4 min 0% B, 10 min 5% B, 13 min 15% B, 13.01 min 20% B until min 15, returning back to 0% at 15.01 min until 18 min. Levels of Kyn were measured at 365 nm absorbance and data analysis was performed with the software Chromeleon v7.2 (ThermoFisher Scientific).

Similar experiments were carried out in wildtype A-673 and SK-N-MC EwS cells after transient transfection with siRNAs. Transfection was performed using Lipofectamine RNAiMAX (Thermo Fisher) on day 0 with either an siRNA targeting EWSR1::FLI1 or a non-targeting control siRNA, followed by re-transfection under the same conditions on day 2, as previously described (Stoll *et al*., 2013; Grünewald *et al*., 2015). Cells were harvested on day 4 after plating.

### NK cell isolation from PBMCs

Buffy coats were obtained from DRK-Blutspendedienst Baden-Württemberg. Peripheral blood mononuclear cells (PBMCs) were isolated by gradient centrifugation with Lymphosep (Lymphocyte Separation Media) (Biowest) and Leucosep tubes (Greiner Bio-One). The PBMC fraction was washed with PBS twice and contaminating red blood cells were lysed using 1x ACK buffer (Elabscience). NK cells were manually isolated with NK cell Isolation Kit, human (Milteny Biotec) and LS columns (Milteny Biotec) following the manufacturer’s protocol. For co-culture experiments, NK cells were resuspended in NK MACS Basal medium (Milteny Biotec), supplemented with 5% human serum Type AB (Fisher Scientific), 1% penicillin/streptomycin and IL-2 (500 U/mL) and IL-15 (140 U/mL) (PeproTech). NK isolation purity percentage (> 90%) was evaluated by flow cytometry (BD FACS Canto II flow cytometry analyzer) and PBMCs, NK cells and NK-depleted fraction cells were stained (1 million cells) with the following antibodies: PE-anti-human CD3, APC anti-human CD56, FITC anti-human CD19, Pacific Blue anti-human CD14 (Biolegend Europe B.V.).

### NK-92 and NK killing assays

To evaluate the killing effect of NK-92 or NK cells against EwS cells, a co-culture was performed and cell viability was evaluated. First, EwS cells (with DOX-inducible KD of *EWSR1::ETS* or *AHR*) were pre-treated with DOX for 72 h (DOX refreshed after 48 h). After 72 h, EwS cells were seeded in 12-well plates (NK-92) or 48-well plates (NK) at equal cell numbers for both DOX conditions and left to attach for 3–4 h. NK-92 or NK cells were then added at a 1:1 ratio to the EwS cells and co-cultured for 24 h. For readout, medium was removed and cells were washed twice with PBS to remove the NK-92 cells. Fresh EwS medium was added and cell viability was determined via resazurin assay as described below.

### Resazurin assay

Resazurin assays were performed to determine cell viability in the drug-response assays following treatment with AHR inhibitors and after co-culture with NK-92 cells. The protocol was followed from already published resazurin colorimetry protocol (Musa and Cidre-Aranaz, 2021) and adapted the volumes accordingly. Resazurin (Sigma-Aldrich, Germany; 1 g/L in PBS) stock solution was diluted 1:10 in PBS to a working stock solution before the assay readout. For NK-92 co-culture experiments, 200 μL resazurin was added to wells containing 500 μL media. For drug response assays, 50 μL resazurin was added to wells containing 200 μL media. Following resazurin addition, plates were incubated for 5 h at 37 °C, and fluorescence readout was performed in the microplate reader GloMax®-Multi Detection System (Promega, USA) with a 560 nm excitation−590 emission filter. Wells without seeded cells (only medium) were use for background subtraction.

### *AHR* KD establishment with pLKO Tet-On construct

*AHR* silencing (transcript ID: Human NM_001621.5) was achieved by cloning the shRNA oligonucleotide sequences into the Tet-pLKO-puro backbone (Addgene #21915, USA). Design of the shRNA sequences were obtained from the database GPP Web Portal. Different criteria for selection were followed: Match position (CDS [coding sequence], 3’UTR [untranslated region]), intrinsic score (<10) and Specificity-Defining Region (SDR) Match (100%). In addition, five shRNA sequences were obtained from the online tool SplashRNA (Pelossof *et al*., 2017). Sequences with a Splash score of 1.5 or higher were selected. The selected sequences were adapted to the Tet-pLKO-puro vector by adding compatible restriction site overhangs of the EcoRI and AgeI enzymes (New England Biolabs, Germany), loop sequence and termination sequence. Details of the cloning protocol including bacteria transformation, single bacteria colony screening, lentivirus production and antibiotic selection has been already established previously by us (Musa, 2021; Orth *et al*., 2022).

### Generation of single cell clones

Lentivirally transduced and puromycin (InvivoGen, Germany) selected cell lines with the DOX-conditional *AHR* KD (bulk level) were seeded in 1:5 collagen coated 96-well plates. To ensure one clone per well, 0.8 cells/well were seeded in a total volume of 200 μL. Medium contained 40% of pre-filtered (0.45 μm) conditional growth medium of the respective cell lines, supplemented with 20% FCS and 1% penicillin/streptomycin. Growth of the single clones was monitored for 1−2 months and Accutase (Sigma-Aldrich, Germany) was used to transfer them consecutively larger plates and flasks for expansion and further KD efficiency testing by qRT-PCR. All single clones were treated with 1 μg/mL DOX (Sigma-Aldrich, Germany) for 96 h, with DOX refreshed every 48 h.

### Proliferation assays

For 2D proliferation assays, the trypan blue (Sigma-Aldrich, Germany) exclusion method was used. Cells were seeded at 2.5−4×10^5^ cells/well for MHH-ES-1/TR/shAHR and 3−5×10^5^ cells/well for TC-106/TR/shAHR in 6-well plates, then exposed to 1 μg/mL DOX for 96 h (DOX refreshed every 48 h) at which point the cell culture supernatant and cells were collected. For counting cells, 10 μL of cell suspension were mixed 1:1 with trypan blue and live and dead cells counted in four quadrants of a hemocytometer. The proliferation relative to control was determined by calculating the average of all DOX− biological independent controls and dividing all cell numbers of all conditions and biological replicates by that average.

### CellTiter-Glo for 3D cell viability assays

For 3D cell viability assays, cells were seeded (MHH-ES-1/TR/shAHR255 and MHH-ES-1/TR/shAHR286: 1,500/well; TC-106/TR/shAHR255 and TC-106/TR/shAHR286: 1,000 cells per well) in 96-well U-bottom low attachment plates (Grenier Bio-one, Germany) to generate a single sphere per well. Spheres were treated with 1 μg/mL DOX for 96 h and DOX was refreshed every 48 h. To determine the effect of AHR silencing on 3D cell viability, a CellTiter-Glo® Cell Viability Luminescent assay (Promega, USA) was performed. After seeding the cells and treating them with DOX for 96 h, spheroids and media blanks (50 μL) were transferred to a white opaque 96-well plate and 50 μL of CellTiter-Glo reagent added per sample. The plate was placed on the shaker for 20 min and luminescence was measured in the GloMax plate reader (1 s integration time). Relative viability compared to the control was calculated.

### Colony formation assay

EwS cell lines were seeded at 2,000 cells/well for MHH-ES-1/TR/shAHR or /shControl, and at 1,000 cells/well for TC-106/TR/shAHR or /shControl. After 24 h of seeding, 1 μg/mL of DOX was added and refreshed every 48 h. In addition, 200 μL of only medium were added to both conditions to take into consideration medium evaporation. Crystal violet (Sigma-Aldrich, Germany) staining was performed after 10−12 d. The size and percentage of area of colonies per condition were calculated with the Fiji software (NHI, USA). Relative clonogenicity was calculated normalizing to the controls.

### Flow cytometric cell cycle analysis

Propidium iodide (PI) (Bio-Legend, USA) staining was performed for cell cycle analysis. MHH-ES-1/TR/shAHR255 (4×10^5^ cells/well) and TC-106/TR/shAHR286 (5×10^5^ cells/well) were seeded in 6-well plates and 1 μg/mL DOX was added for 96 h. On the day of harvesting, medium was removed from the cells and were washed with ice-cold PBS (ThermoFisher Scientific, USA). Cells were trypsinzed and centrifuged down. After pellet collection cells were washed again with PBS and then fixated with 1 mL of cold 70% ethanol (dropwise). To perform the staining, samples were incubated in the following master mix: 470 μL of flow cytometry staining buffer, 20 μL of PI solution and 10 μL of RNAase (ThermoFisher Scientific, USA). Incubation was performed for 1 h in the dark. Samples acquisition was performed on a BD FACSCanto flow cytometer (BD Biosciences, USA) with 0.5−1×10^5^ events at medium flow rate per sample. Data analysis was performed using the FlowJo v.10.10 software (BD Biosciences, USA), with the percentage of cells present in different cell cycles (sub G1/G0, G1/G0, S and G2/M) determined using the Watson model.

### Mouse experiments

All mouse experiments were conducted with the approval of the government of North Baden as the responsible legal authority. The study adhered to the ARRIVE guidelines, European Community and GV-SOLAS recommendations (86/609/EEC), and United Kingdom Coordinating Committee on Cancer Research (UKCCCR) guidelines for the welfare and use of animals in cancer research. Animals were housed at the central DKFZ animal facility under Specific Opportunist Pathogen-Free (SOPF). Animals had *ad libitum* access to food and water and were housed in individually ventilated cages. Daily assessments of animal well-being were carried out by certified animal caretakers. Mice subcutaneous (s.c.) xenografts and orthotopic xenografts for metastasis assessment were performed as previously described (Cidre-Aranaz and Ohmura, 2021; Cidre-Aranaz *et al*., 2022). In brief, for s.c. experiments, 3×10^6^ EwS cells (MHH-ES-1/TR/shAHR255 or TC-106/TR/shAHR286) resuspended in 100 μL of a 1:1 mix of Geltrex:PBS (ThermoFisher Scientific, USA) were inoculated subcutaneously into the right flank of each NSG mouse. For orthotopic experiments, 2×10^5^ MHH-ES-1/TR/shAHR255 cells were resuspended in 10 μL HBSS 1× and inoculated into the right tibial plateau of each NSG mouse. Once tumors were palpable (s.c.) or the day after injection (orthotopic), mice were randomized and received treatment in the drinking water according to their grouping: control (1.75% sucrose; Sigma-Aldrich, Germany) (DOX−) or treatment (2 mg/mL DOX in 5% sucrose, DOX+). Drinking water was refreshed every 48 h. Primary tumor at the s.c. or orthotopic location were measured every second day with a caliper and volumes were calculated using the following formula: V=L×W^2^/2, where V is the tumor volume, L the largest diameter and W the smallest diameter. For both s.c. and orthotopic experiments, mice were sacrificed by cervical dislocation at the experimental endpoint or earlier if humane endpoints were reached (body weight loss of 20%, apathy, piloerection, self-isolation, aggression as a sign of pain, self-mutilation, motor abnormalities, as well as any other unphysiological or abnormal body posture, breathing difficulties, maximum tumor size of 15 mm in any direction, an ulcerating tumor or exhibited signs of limping at the injected leg).

### DNA isolation

DNA concentration and purity were determined using a NanoDrop One/One UV-Vis spectrophotometer (ThermoFisher Scientific, USA). Genomic DNA for SNP or STR profiling was extracted from fresh or frozen culture cells using the NucleoSpin® Tissue kit as per manufacturer’s protocol. Samples for profiling were submitted with a final concentration of 15 ng/μL in 100 μL. DNA from transformed bacteria was isolated using ZymoPURE II Plasmid prep kits as per manufacturer’s protocols (Zymo Research, USA). To maximize the amount of DNA isolated, the elution buffer was pre-heated at 70 °C and added right before the elution to the column. To isolate DNA from the agarose gel, NucleoSpin Gel^TM^ and PCR Clean-up kits (Macherey-Nagel, Germany) were used according to the manufacturer’s protocol.

### Immunoblotting

*AHR* silencing was confirmed at the protein level. All single clones were treated with 1 μg/mL DOX for 96 h and DOX was refreshed every 48 h. Cells were collected following 96 h DOX treatment and lysed in RIPA buffer containing phosphatase inhibitor and protease inhibitor. Protein was quantified by Pierce BCA Protein Assay Kit (ThermoFisher Scientific, USA) as per manufacture’s protocol. Samples (20 μg whole cell lysate with 6× Laemmli loading buffer in a total volume of 20 μL) were run on 10% Tris-HCl gels (Bio-Rad, Hercules, CA, USA). Meanwhile, PVDF membrane was activated in 100% methanol the equilibrated in transfer buffer. Proteins were transferred to the membrane at 1.0A for 7 min using the Turbo transfer system (Bio-Rad). Membrane was rinsed in TBS-T then blocked in 5% milk/TBS-T for 1 h at room temperature. Membranes were washed three times for 5 min in TBS-T then incubated with 1:1,000 anti-AHR (D5S6H clone, #83200, Cell Signaling Technologies, Danvers, MA, USA) or 1:1,000 anti-GAPDH (14C10 clone, #2118S, Cell Signaling Technologies) antibodies in 5% milk/TBS-T, overnight at 4 °C. Membranes were then washed three times for 5 min with TBS-T and then incubated with HRP-conjugated goat anti-rabbit secondary (1:2,000, 5% milk/TBS-T, 2 h, room temperature; #31460, ThermoFisher Scientific). Then, the membrane was washed three times for 5 min in TBS-T and incubated with WesternBright chemiluminescence substrate (Biozym, Germany) for 2 min. Proteins were visualized using the FusionFX Western Blot & Chemi imaging system (Vilber, France).

### Drug-response assays

MHH-ES-1 and TC-106 cells were seeded (4,000 cells/well) in 96-well plates and treated with the AHR inhibitor GNF-351 (MedChemExpress, USA; 200 mM in DMSO) for 72 h. Resazurin assay was performed to determined cell viability as described above. CFA assays were performed in a dose-dependent matter by treating EwS cells with GNF-351 for 72 h. Staining of colonies and analysis was performed as described before.

### Statistics

Statistical analysis of both *in vitro* and *in vivo* experiments was performed using GraphPad PRISM 10.1.2 (GraphPad Software, USA). Data were presented as mean ± standard error of the mean (SEM), and if not otherwise specified one- or two-sided Mann-Whitney test was performed.

---

**Supplementary Figure 1.**
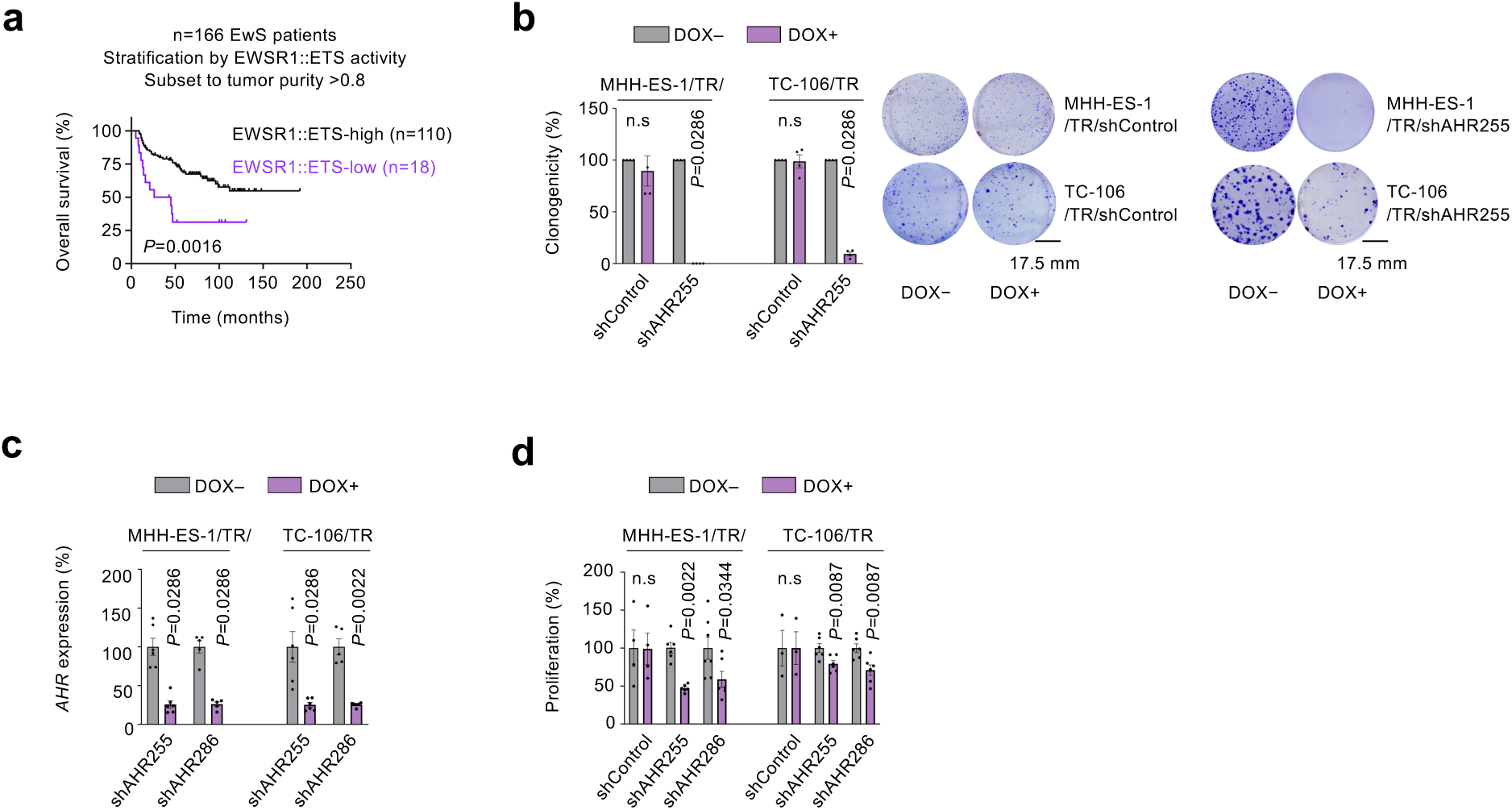
Low EWSR1::ETS activity in EwS is associated with poor clinical outcome, and silencing of AHR leads to decreased EwS cell proliferation and clonogenicity. **a)** Kaplan-Meier survival analysis of n=128 patients stratified by EWSR1::ETS-activity and with an inferred tumor purity of >80% (log-rank test). **b)** Analysis of relative clonogenicity of indicated cell lines harboring either a DOX-inducible control shRNA (shControl) or shRNA against AHR (bulk level) following 10–12 d of DOX-treatment (left) and representative colony images (right). Horizontal bars represent the mean and whiskers the SEM. n=4 biologically independent experiments. Two-sided Mann-Whitney test. **c)** Relative *AHR* mRNA expression following 96 h DOX treatment in MHH-ES-1/TR/shAHR and TC-106/TR/shAHR cells (bulk level). Horizontal bars represent the mean and whiskers the SEM. n=5−6 biologically independent experiments. Two-sided Mann-Whitney test. **d)** Analysis of proliferation in MHH-ES-1/TR/shAHR and TC-106/TR/shAHR after 96 h of DOX-induced independent shRNAs targeting AHR (255 and 286) or shControls. Counts were performed via trypan blue exclusion method. Horizontal bars represent the mean and whiskers the SEM. n=3−7 biologically independent experiments. Two-sided Mann-Whitney test.

## Notes

https://www.ncbi.nlm.nih.gov/geo

https://github.com/ahmedasadik/AffyGEx

### Summary of Updates

New data has been added to Figure 1

